# Application of nanopore sequencing to identify antimicrobial resistance genes, mobile genetic elements and virulence factors in clinical isolates

**DOI:** 10.1101/2023.10.25.563907

**Authors:** Rachel Kimani, Sebastian Musundi, Patrick Wakaba, David Mbogo, Suliman Essuman, Bernard N. Kanoi, Jesse Gitaka

## Abstract

The global health challenge posed by the emergence of antibiotic resistance pathogen is further exacerbated in African countries by the indiscriminate use of antibiotics, poor surveillance and lack of stewardship programs. To address this issue, we employed the Oxford Nanopore Technologies (ONT) to sequence 17 clinical isolates from a referral hospital in Kenya. Our comprehensive bioinformatics approach facilitated the assembly, identification of sequence types and prediction of antimicrobial resistance genes, mobile genetic elements (plasmids and integrons) and virulence genes. Of the 17 isolates, five were *A. baumannii*, four *E. coli*, three *S. haemolyticus*, three were *E. cloacae,* while *S. aureus* and *E. faecalis* were single isolates. For the detection of AMR genes, *A. baumannii* isolates harbored genes such as *blaOXA-23* which mediates resistance to carbapenems, *E. coli* and *E. cloacae* carried *blaCTX-M-15* which confers resistance to cephalosporins and *S. haemolyticus* harbored *blaZ,* responsible for resistance against *penicillins*. *S. aureus* co-haboured *mecA* and *blaZ genes.* In addition,, various other different AMR genes to chloramphenicol, macrolides, aminoglycosides, tetracycline were also observed. For plasmid replicons, *E. coli* carried the most number of plasmids and shared ColRNAI_1 and IncFIB(pB171)_1_pB171 with *A. baumannii* and IncR_1 with *E. cloacae.* Many genes encoding various virulence factors including *fimA-I* and ompA*, senB* were identified in *E. coli, hlgA-C* and *hla/hly, hlb, hld* in *S. aureus* and *efaA*, *ebpA-C* in *E. faecalis.* In conclusion, most isolates contained a combination of different AMR genes harbored in plasmids and integrons and virulence genes. This study provides significant information on genetic determinants of antibiotic resistant pathogens in clinical isolates and could assist in developing strategies that improve patient treatment.

**Author Summary:** Antimicrobial resistance remains a major health challenge across the globe. The continued misuse and lack of proper monitoring has worsened the problem of antibiotic resistant infections. In this study, we sought to use nanopore sequencing to identify antibiotic resistance genes, mobile genetic elements and virulence factors from clinical isolates which showed resistance against commonly used antibiotics. We found the presence of resistance genes to multiple different antibiotics including beta-lactams, macrolides, tetracycline and aminoglycosides across multiple bacterial species. We also identified plasmid replicons and class I integrons which facilitate the spread of antimicrobial resistant genes. Furthermore, several virulence factors that help resistant bacteria to survive were identified. Overall, this study highlights the widespread issue of antibiotic resistance, factors contributing to its persistence in clinical isolates and utility of nanopore sequencing for monitoring genetic determinants of antimicrobial resistance.

## Introduction

Over the past 70 years, treatment and control of bacterial diseases has greatly relied on use of antibiotics [1]. However, the rise in antimicrobial resistance (AMR) in the past decades has emerged as a major public health challenge globally[2]. With the escalating spread of multi-drug resistant bacteria, widely used combinations of empirical antibiotic treatment regimens are being challenged [3]. In particular, infections caused by methicillin resistant *Staphylococcus aureus* (MRSA), extended-spectrum beta-lactamases (ESBLs) producing gram negative bacteria and Carbapenem-resistant Enterobacterales (CREs), poses a major health concern. These bacteria are not only resistant to all penicillin and third generation cephalosporins, but also frequently express resistance against carbapenem. As a result, treatment failure may occur leading to longer hospital stays and at worst death.

Early identification and routine epidemiological surveillance of AMR is necessary so as to provide informed use of antibiotics and hence prevent further spread of AMR [4]. Phenotypic identification of antimicrobial susceptibility profiles of bacterial isolates is commonly performed by microbiological procedures based on isolation and growth of pure cultures of individual bacterial isolates in the presence of an antimicrobial agent. These methods suffer from long turn-around time and the methods only allow assessment of resistance to known antibiotics. Whole genome sequencing offers several advantages over traditional culture-based approaches including its ability to accurately identify strains and comprehensively identify AMR genes, plasmids, and virulence factors [5]. Already the use of WGS to identify and characterize antimicrobial resistant bacterial strains has significantly increased over the past years [6, 7]. Nonetheless, several limitations, key among them cost hinder the adoption and deployment of WGS in resource constrained settings.

In April 2014, the Oxford Nanopore Technologies (ONT) launched a portable long-read nanopore sequencing MinION device that has comparable performance to standard short-read sequencing platforms. ONT sequencing offers unique advantages to short-read sequencing including lower equipment costs, reduced turnaround times from days to hours, real-time base-calling, enrichment of samples via adaptive sampling and high portability [8, 9]. To date, ONT MinION sequencer has been widely applied in research to detect bacteria, parasites and viruses in both clinical and environmental samples [10–12]. The MinION device has been used in detecting bacterial pathogens including *Salmonella typhirium* [13], *Salmonella enterica* and MRSA [14]. In this study, we performed whole genome sequencing for 17 clinical samples that harbored isolates such as *A. baumannii*, *E. cloacae, E. faecalis* and *Staphylococcus spp* using ONT to screen for genes that confer resistance against commonly used antibiotics. We identified antimicrobial genetic determinants such as plasmids, integrons, and virulence genes harbored by the bacterial isolates.

## Results

### Genome Assembly Quality and Completeness

The total number of contigs from the draft genomes ranged from 2 to 61, with the largest contig being approximately 4,948,395 bases. The N50 value ranged from 236,330 to 4,948,395, while the average GC% content ranged from 32.78% to 55.89 (S1 Table). Genome completeness before polishing by BUSCO using the proteobacteria database ranged from 37.9% to 99% (S1 Fig). After four rounds of polishing using Racon followed by Medaka and Homopolish, the BUSCO completeness score ranged from 44.7% to 100% (S2 Fig).

### Sequence Types

MLST analysis of the 17 isolates revealed different bacterial species with varying sequence types. Of the 17 isolates, 5 were *A. baumannii,* 4 were *E. coli,* 3 were *S. haemolyticus,* 3 were *E. cloacae,* while *S. aureus* and *E. faecalis* were single isolates (S2 Table). One sequence type in *A. baumannii* isolates could be identified as ST1. The other *A. baumannii* isolates showed high similarity ∼98% and coverage ∼99% with other sequence types (S2 Table). For *E. coli* isolates, ST53 was present in 3/4 isolates, with one isolate showing close similarity to ST-110,143 and 121 due to 4 gaps in fumC. The *S. haemolyticus* isolates were of ST30 and ST1, while the remaining isolate showed close similarity to ST65,15 or ST1. The three *E. cloacae* isolates were ST88, ST98, and ST78, *S. aureus* isolate was ST6, and the *E. faecalis* isolate was ST538 (S2 Table).

### Antimicrobial resistance Genes

Identification of AMR genes determinants from whole genome sequencing data revealed different AMR determinants for various bacterial species. All 17 sequenced bacterial isolates carried a number of antibiotic resistant genes. Notably, *A. baumannii* isolates carried the most ARG’s and this was also observed phenotypically in terms of resistance to more than three different antibiotics class. *A. baumannii* isolates carried beta-lactamase resistance genes including *blaOXA-23, blaOXA-276, blaOXA-304, blaOXA-317, blaOXA-507*, and *blaOXA-69*. Additionally, we found other beta-lactamase resistance genes, namely; *blaADC-134, blaADC-174, blaADC-5, blaADC-73, blaCMY-127,* and *blaPER-7*. Moreover, some isolates showed resistance to Macrolide, Lincosamide, Streptrogramins (MLS) (*mph(E)* and *msr(E))*, Amphenicols (*Cat1* and *cmlA5I),* tetracycline (*tet(B)* and *tet(X),* aminoglycosides (*armA, aph(6)-Id* and *aph(3’’)-Ib)*, rifamycin (arr-2), quinolone (*qnrB29)* and sulfonamides (*sulf1* and *sul2*). Most *A. baumannii* isolates (4/5) contained *ant (3’’)-IIa,* which confer resistance against aminoglycosides spectinomycin and streptomycin (Fig 1A).

**Fig 1.**
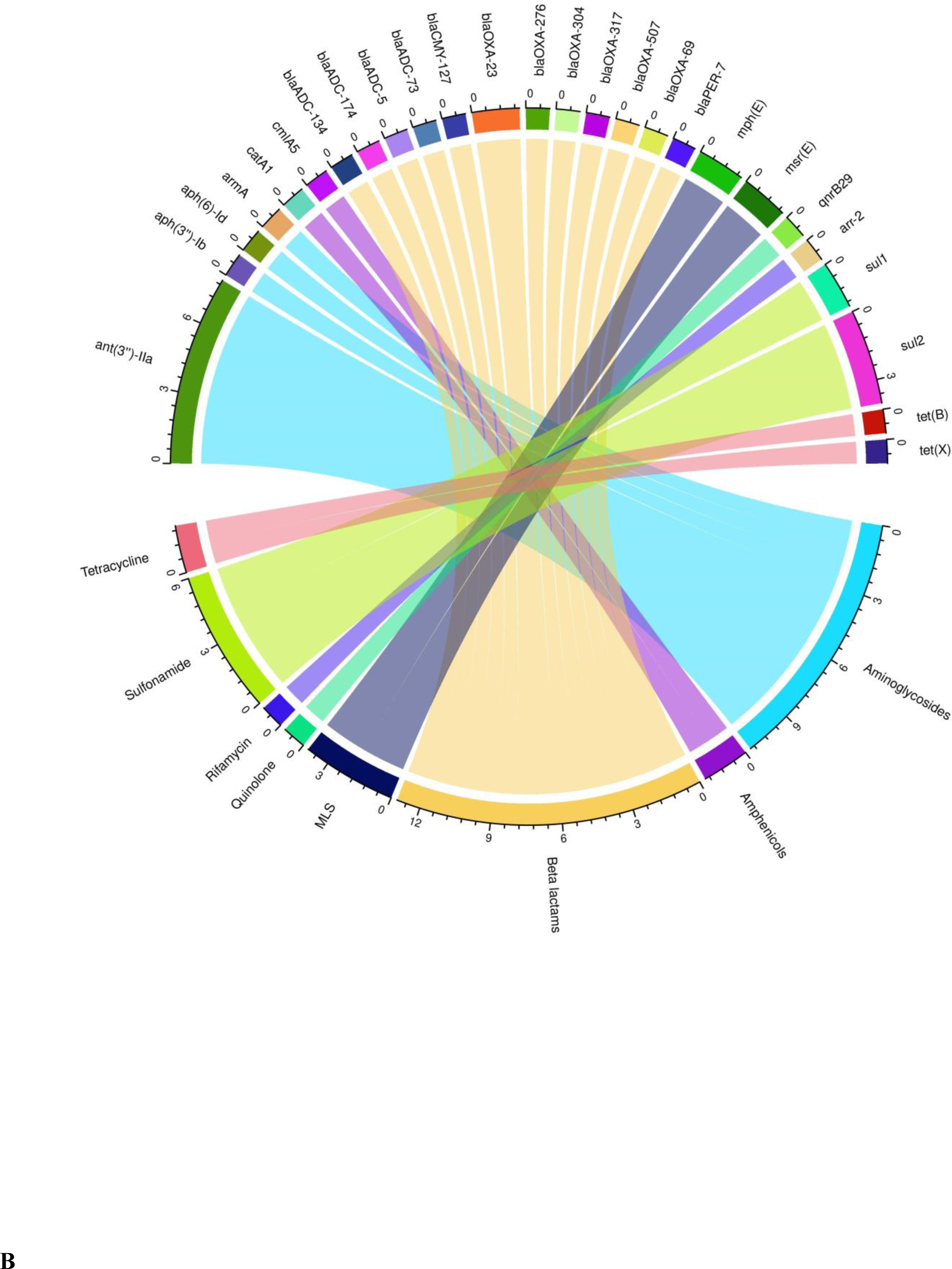
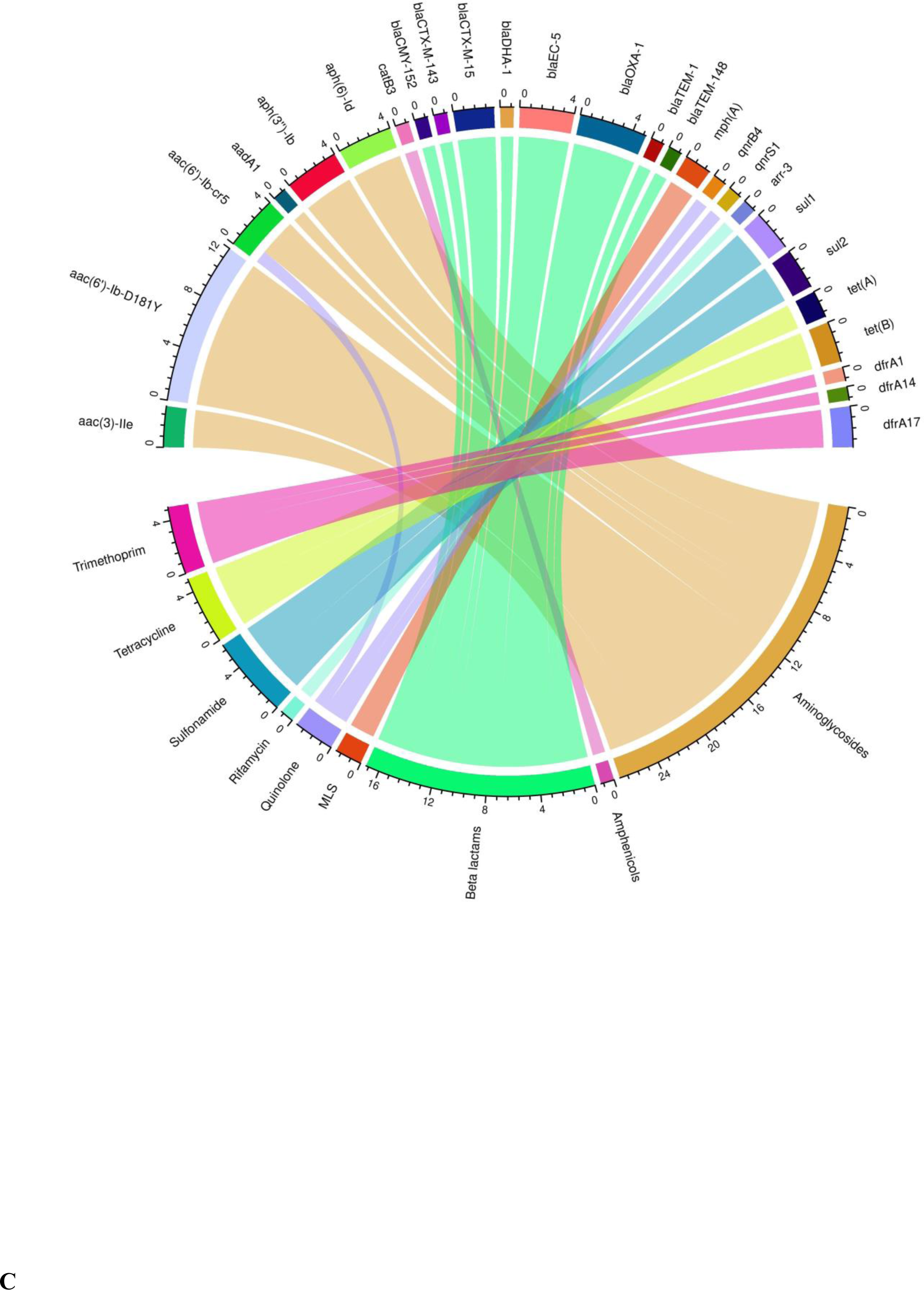
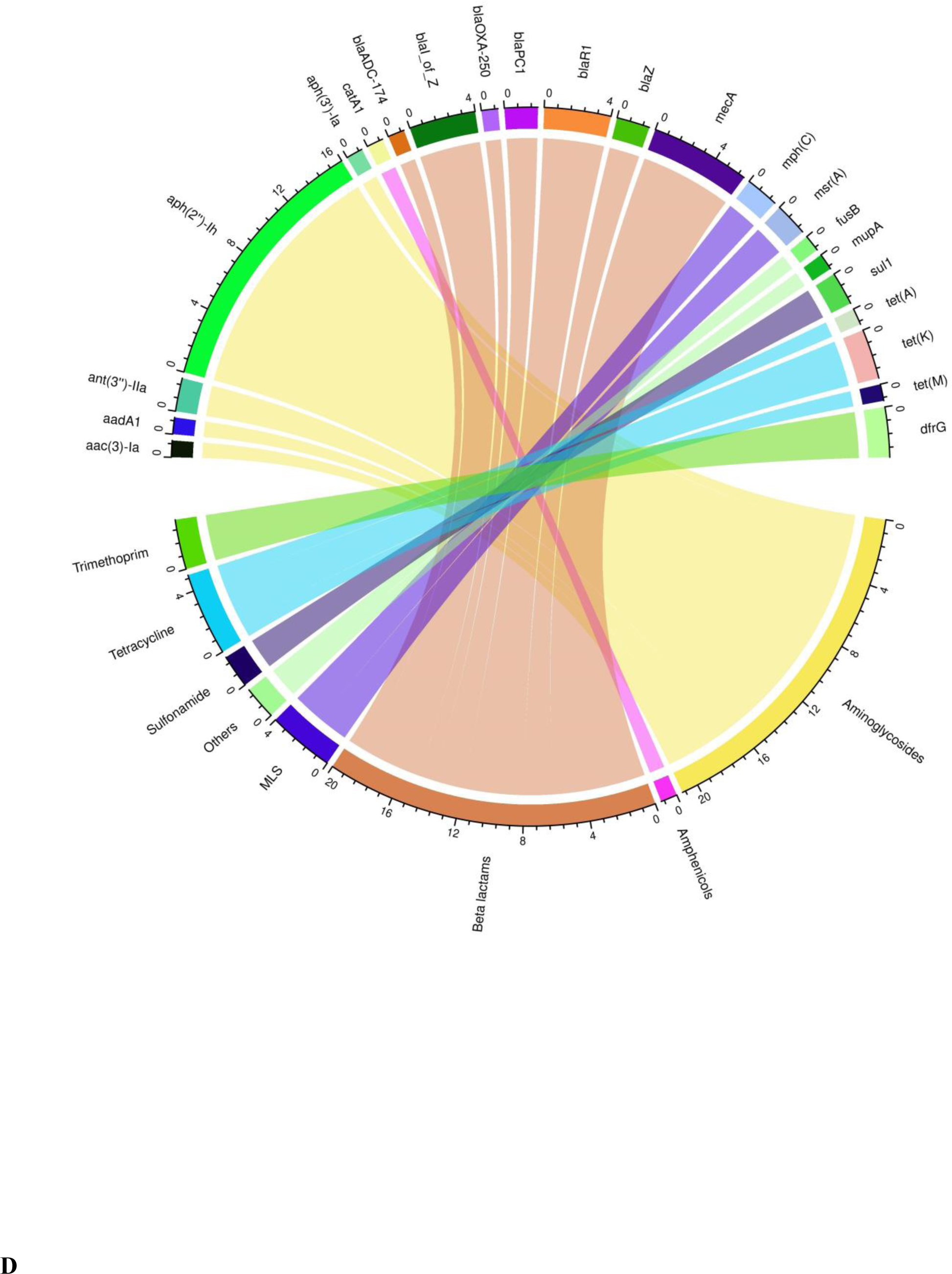
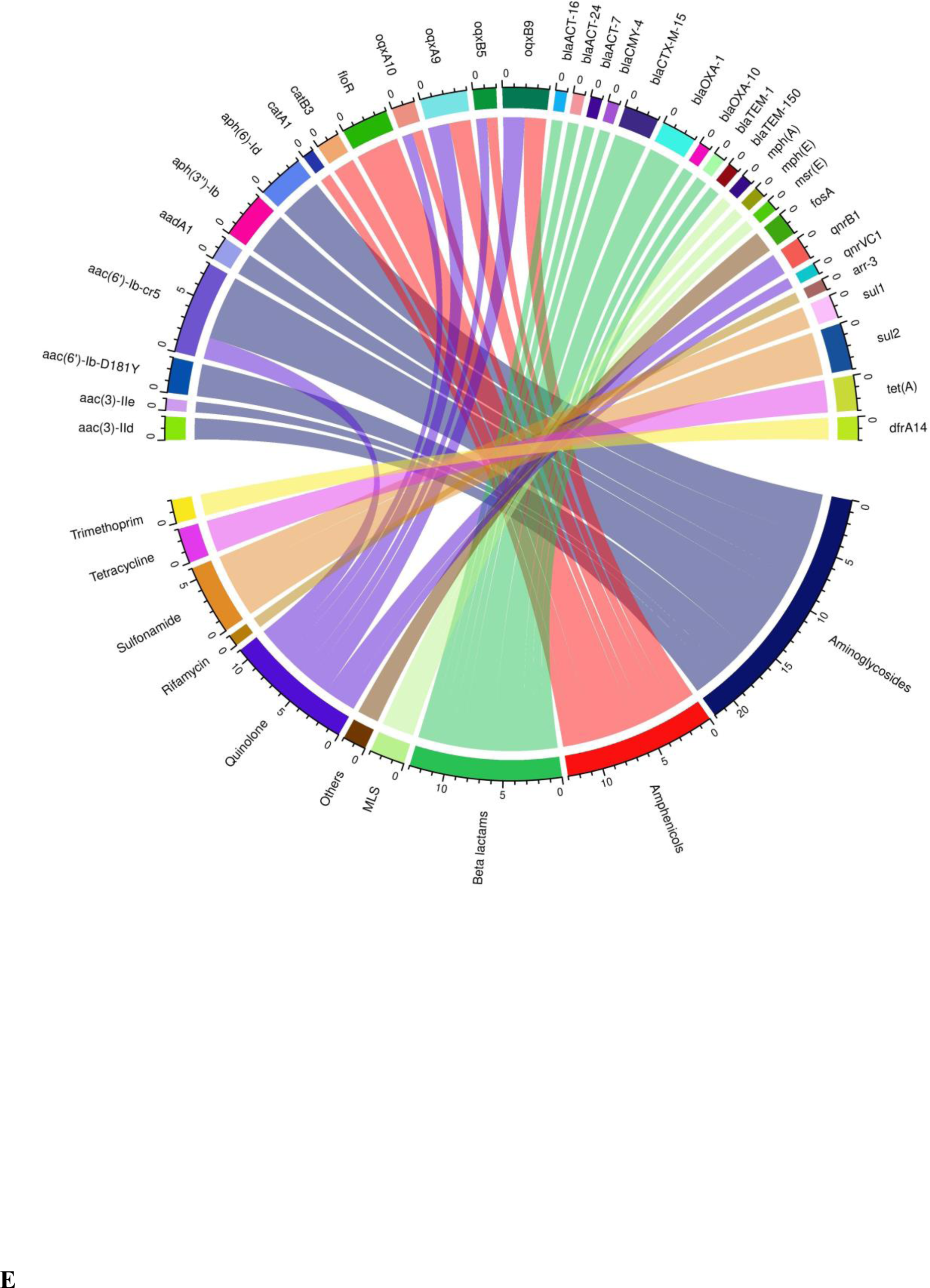
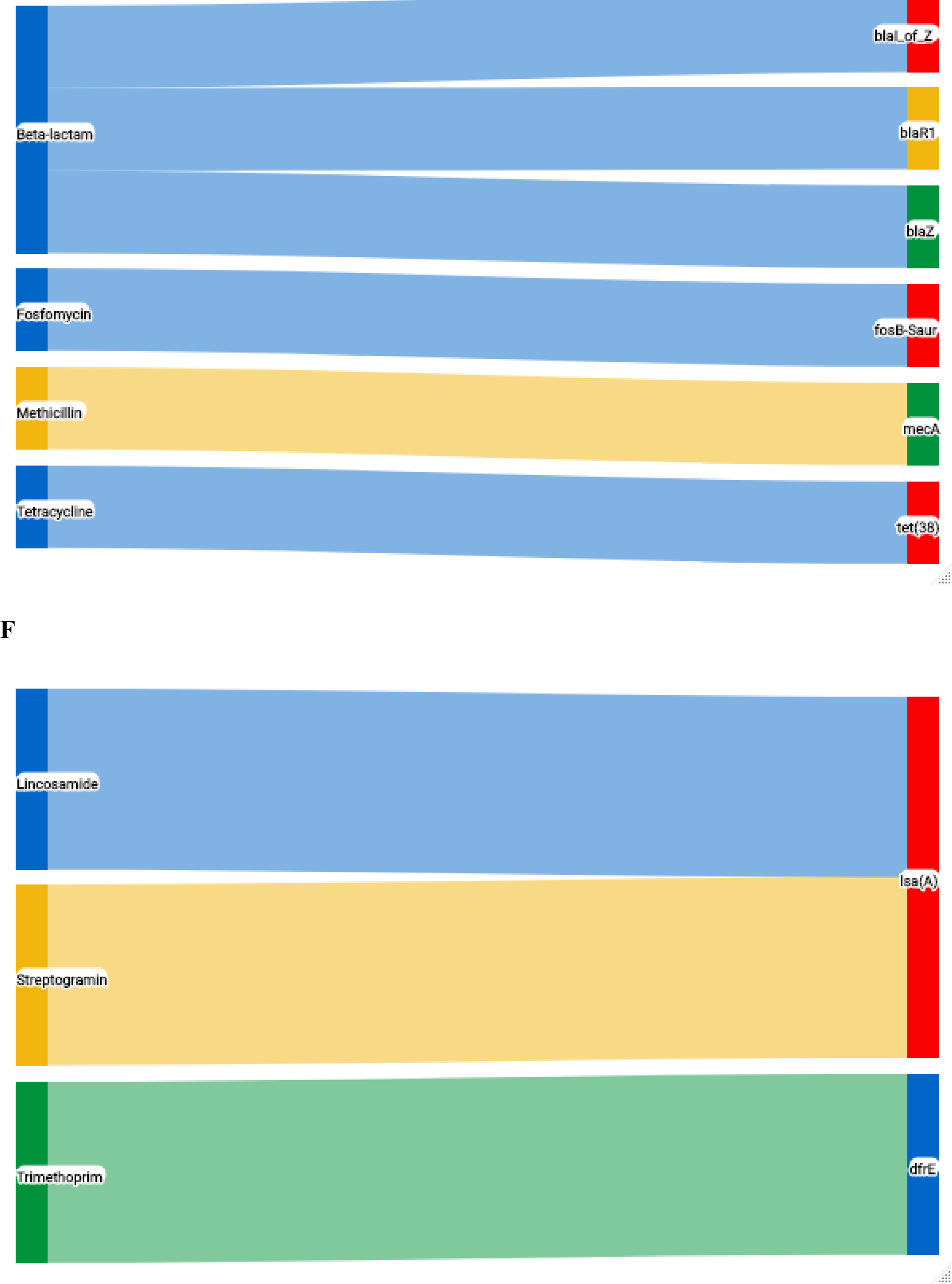
Predicted antimicrobial resistance genes and their antibiotic class. Circos plot linking the number of antimicrobial resistance genes and class in **A**) *A. baumannii* **B**) *E. coli* **C**) *S. haemolyticus* **D**) *E. cloacae.* Sankey chart linking the antimicrobial resistance gene and class in ***E*)** *S. aureus* **F**). *E. faecalis*

In the four *E. coli* isolates, we identified beta-lactam resistance genes *blaTEM-1*, *blaTEM-148*, *blaCTX-M-15,* and *blaOXA-1*. Other beta lactam resistance genes were also identified including *blaEC-5, blaDHA-1, blaCTX-M-143,* and blaCMY-152. Additionally, resistance genes against amphenicol chloramphenicol (*catB3*), aminoglycosides gentamicin (*aac(3)-IIe*), kanamycin(*aac(6’)-Ib-cr5, aac(6’)-Ib-D181Y*), streptomycin(*aph(3’’)-Ib, aph(6)-Id, aadA1*), tobramycin (*aac(6’)-Ib-D181Y, aac(6’)-Ib-cr5*), as well as MLS (*mph(A)*), quinolones (*qnrB4, qnrS1*), tetracycline (*tet(A), tet(B)*), trimethoprim(*dfrA14, dfrA14, dfrA17*), rifamycin (*arr-3*), and sulfonamides (*sul1* and *sul2*) were present in *E. coli* isolates (Fig 1B). Similar to *A. baumannii,* most *E. coli* isolates exhibited multidrug resistance.

A total of 26 different antimicrobial resistance genes were observed among the four isolates of *S. haemolyticus.* The detected genes included those encoding resistance against beta-lactam including *blaI_of_Z, blaR1, blaPC1, blaZ, blaOXA-250* and *blaADC-174* and *MecA*. Other resistance genes present in the *S. haemolyticus* isolates conferred resistance against aminoglycosides amikacin (*aph(2’’)-Ih*), kanamycin (*aph(2’’)-Ih, aph(3’)-Ia*), gentamicin (*aph(2’’)-Ih, aac(3)-Ia*), streptomycin (*aadA1 and ant(3’’)-IIa*), and spectinomycin (*ant(3’’)-IIa*). In addition, resistance genes were also detected against amphenicols (*catA1*), fusidic acid (*fusB*), MLS (*mph(C), msr(A*)), tetracycline (tet*(K), tet(A),* and *tet(M)*), and trimethoprim (*dfrG*) (Fig 1C).

The *E. cloacae* isolates demonstrated a diverse array of 44 different ARG. They included resistance genes against beta-lactam *blaCTX-M-15, blaOXA-1, blaACT-16, blaACT-24, blaACT-7, blaCMY-4, blaOXA-10, blaTEM-1*, and *blaTEM-150*. Additionally, resistance genes that were observed in *E. cloacae* isolate conferred resistance against aminoglycosides amikacin (*aac(6’)-Ib-cr5*, and *aac(6’)-Ib-D181Y*), kanamycin(*aac(6’)-Ib-cr5, aac(6’)-Ib-D181Y*), streptomycin(*aph(3’’)-Ib, aph(6)-Id, aadA1*), tobramycin *(aac(6’)-Ib-cr5, aac(6’)-Ib-D181*Y) and gentamicin(*aac(3)-IId, aac(3)-IIe*). Furthermore, resistance genes against Amphenicols chloramphenicol (*catB3, floR, catA1*), phenicols (*oqxA9, oqxB9, oqxA10, oqxB5*), florfenicol(*floR*), fosfomycin(*fosA*), as well as MLS (*mph(A), mph(E), msr(E)*), quinolones (*aac(6’)-Ib-cr5, oqxA9, oqxB9, qnrB1, oqxA10, oqxB5, qnrVC1),* rifampicin (*arr-3*), sulfonamides (*sul2, sul1),* tetracycline*(tet(A)),* and trimethoprim (*dfrA14*) were also identified (Fig 1D). The single *S. aureus* isolates carried resistance genes against methicillin (MecA), tetracycline(*tet(38*)), fosfomycin(*fosB-Saur*), as well as other beta-lactam resistance genes *blaR1, blaZ* and *blaI_of_Z* (Fig 1E). *E. faecalis* isolate carried the resistance gene *dfrE,* resulting in resistance against trimethoprim and *lsa(A),* which confers resistance to streptogramin and lincosamide (Fig 1F).

### Plasmids and Integrons

Plasmid replicons were identified in 8 out of 17 isolates spanning different bacterial species. Specifically, plasmid replicons were found in *A. baumannii* (1/5), *E. coli* (3/4), *E. cloacae* (3/3), *S. aureus* (1/1), and *S. haemolyticus (3/3).* IncF plasmids were identified as the most common replicon type and were present in *E. coli, E. cloacae* and *A. baumannii* isolates. Other plasmids types present in *E. coli* isolates included Col, IncI, IncR, and RepA. In *A. baumannii,* the other identified replicon type was Col while pENTAS02, IncR, IncA replicon types were present in *E. cloacae.* The replicon types repUS43, rep7a and rep39 were present in the *S. haemolyticus* isolates while rep5a and rep16 were present in *S. aureus* (Fig 4A). Interestingly, some plasmids were shared across different bacterial species. For example, ColRNAI_1 and IncFIB(pB171)_1_pB171 replicons were present in both *A. baumannii,* and *E. coli* isolates, while IncR_1 was present in *E. coli* and *E. cloacae* (Fig 2A). In *E. coli*, plasmids Col156_1, IncFIA_1, IncFIB(AP001918)_1, and IncI1_1_Alpha were present in all three isolates (Fig 3). In addition, the plasmids RepA_1_pKPC-CAV1321 and IncFII(Yp)_1_Yersenia were present in individual isolates of *E. coli* and *A. baumannii,* respectively. Notable plasmids identified in *E. cloacae* included IncA/C2_1, IncFIA(HI1)_1_HI1, IncFIB(K)_1_Kpn3, IncFIB(pECLA)_1_pECLA, IncFIB(pHCM2)_1_pHCM2 and pENTAS02_1. The plasmids rep39_1_repA (SAP110A) and rep7a_16_repC (Cassette) were present in 2 *S. haemolyticus* isolates, while repUS43_1_CDS12738(DOp1) was present in a single isolate. The *S. aureus* isolates contained the plasmid replicons rep5a_1_repSAP001(pN315) and rep16_3_rep(pSaa6159) (Fig 2B).

**Fig 2.**
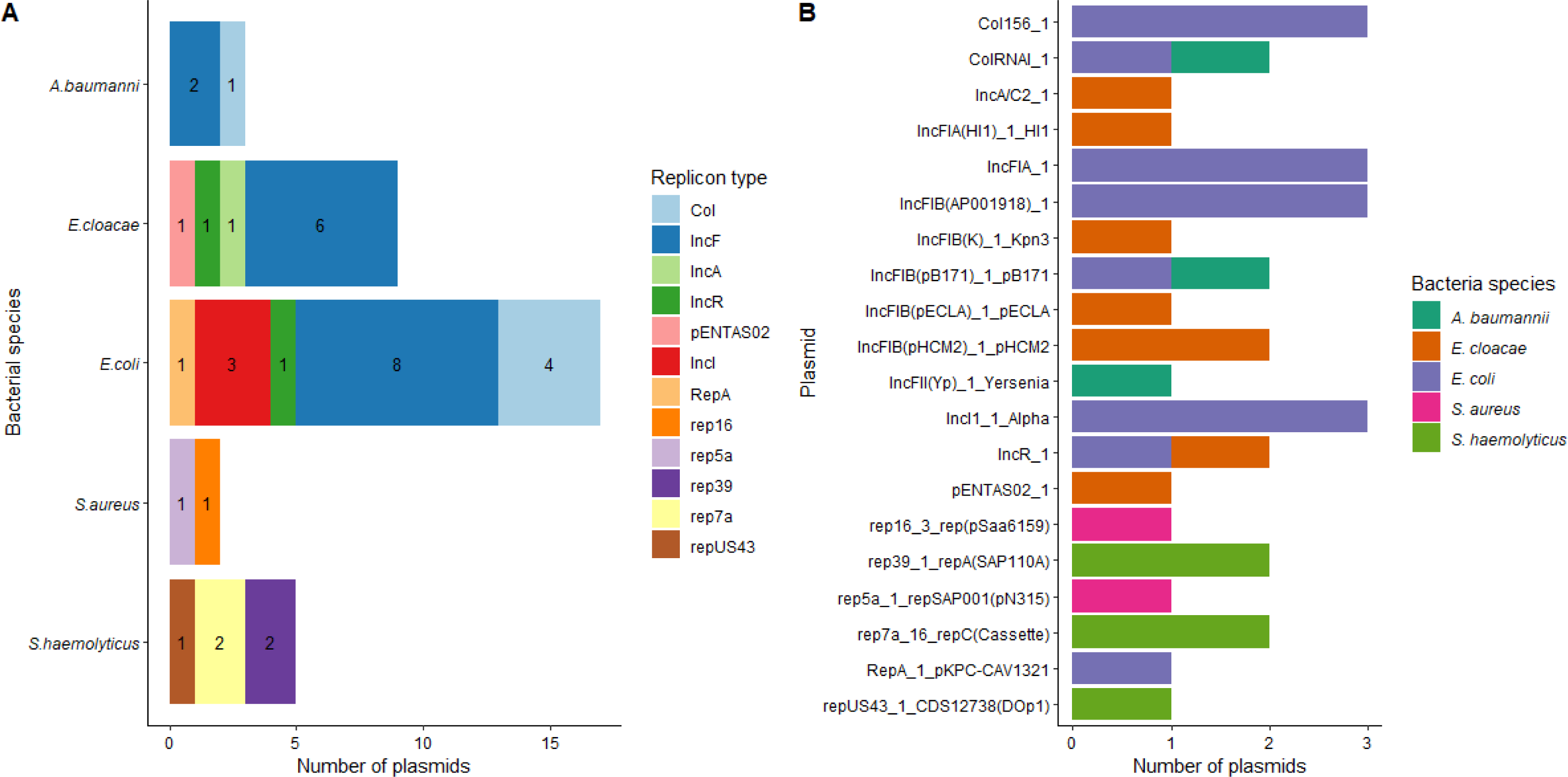
Plasmids identified in different clinical isolates. **A)** Species distribution of individual plasmid types. **B**). Number of individual plasmid types identified across different bacterial species.

**Fig 3.**
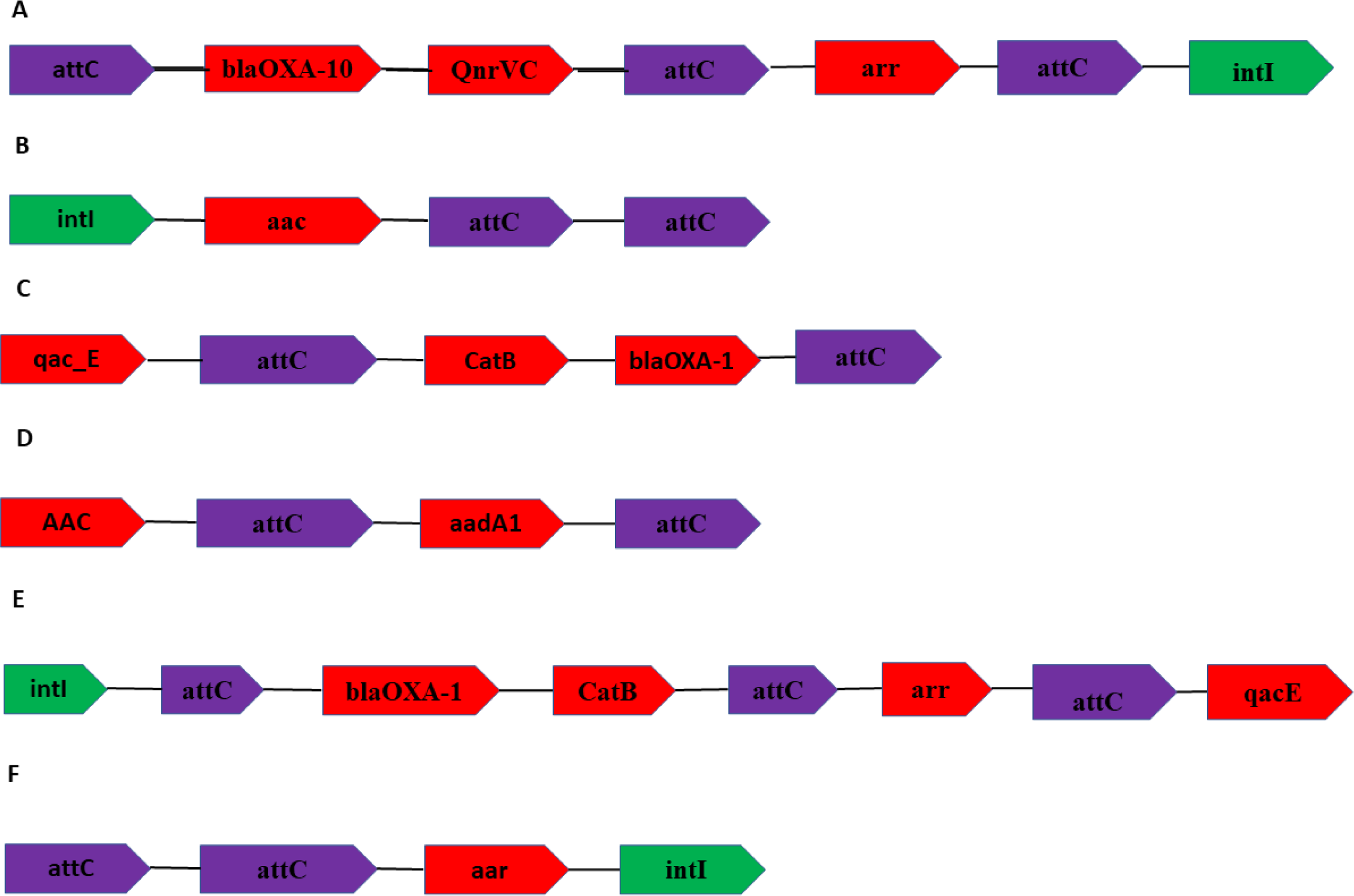
Distribution of different integrons identified in bacterial isolates. The integrons have been identified from **A-C)** *E. cloacae*, **D)** *S. haemolyticus*, **E)** *E. coli*, **F)** A. *baumannii*. Class I integrons consists of three parts including IntI (Integron integrase), enzyme responsible for site specific recombination, attC site which are recognized and recombined by IntI to incorporate new gene cassettes. The attC sites are essential for class I integron ability to capture and express various gene cassettes including those that confer antibiotic resistance to beta-lactamases (*blaOXA-10, blaOXA-1*), quinolones (*QnrVc*), aminoglycosides (*arr, aac, aadA1*), disinfectants and antispetics(*qac_E*), and chloramphenicol (*CatB*)

**Fig 4.**
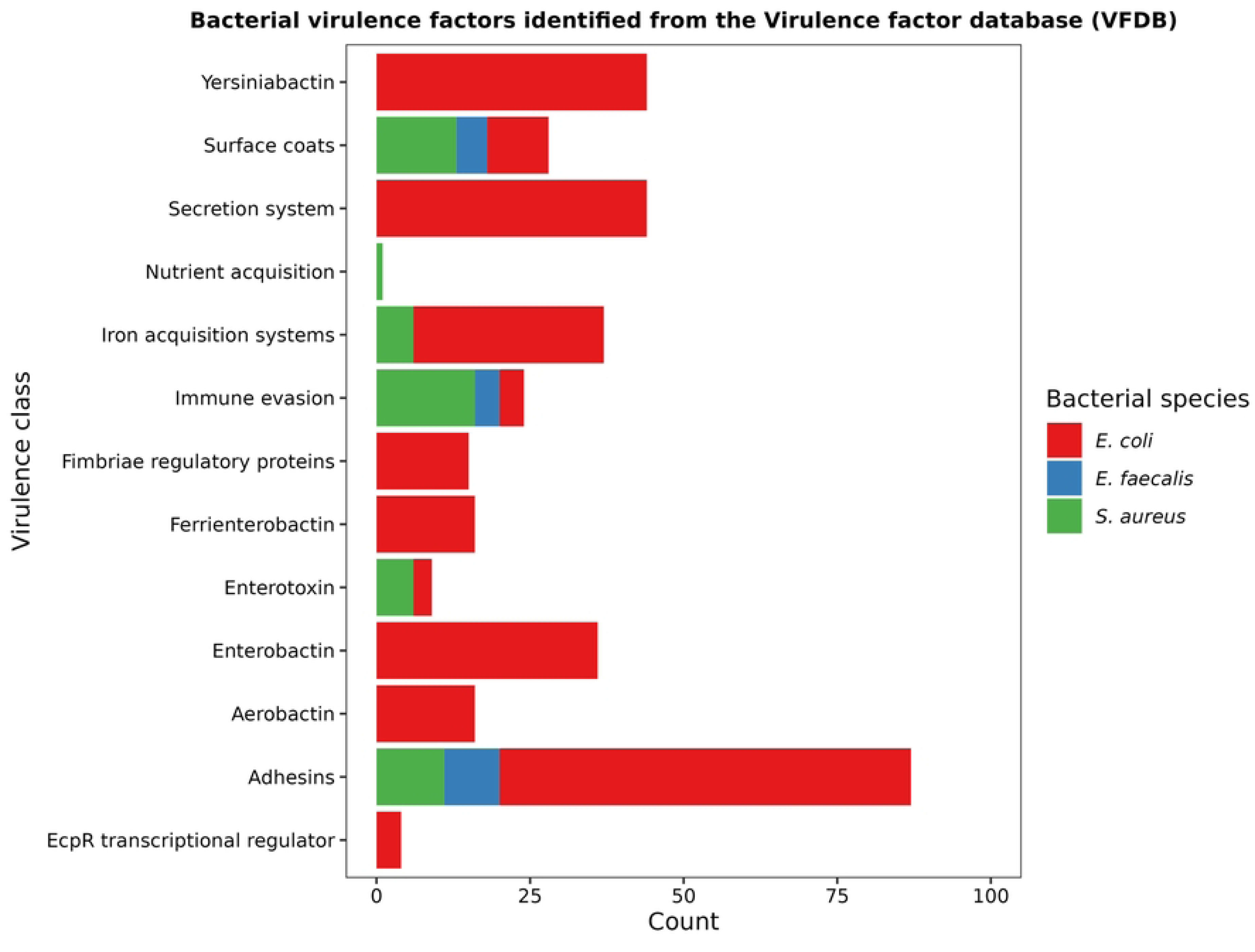
Virulence gene classes identified in *E. faecalis*, *S. aureus*, and *E. coli* isolates.

For integrons, the *E. cloacae* isolates contained three different integron class 1 integron, which contained various AMR genes, including *blaOXA-1, qnrVC, arr, QacE, catB, blaOXA-10* which conferred resistance to beta-lactams, quinolones, rifampicin, chloramphenicol, and quaternary compounds (Fig 3A-C). *S. haemolyticus* isolate carried a class 1 integron containing the resistance genes *aac* and *aadA1* (Fig 3D), associated with resistance to aminoglycoside, streptomycin, and spectinomycin, respectively. Furthermore, the *E. coli* isolate class 1 integron carried the genes *blaOXA-1, catB, arr,* and *qacE,* conferring resistance to beta-lactams, chloramphenicol, rifampicin, and quaternary compounds, respectively (Fig 3E). Lastly, the *A. baumannii* isolate contained an integron that carried the resistance gene *aar,* responsible for rifampicin resistance (Fig 3F).

### Virulence Genes

Virulence-associated genes associated with adherence, immune evasion, secretion systems, iron acquisition, iron acquisition systems and siderophores, nutrient acquisition, secretion systems, and surface coats were identified across *E. coli*, *E. faecalis,* and *S. aureus* isolates. The majority of virulence genes were detected in *E. coli* isolates (n=79), followed by *S. aureus* (n=51) and *E. Faecalis* (n=18)(Fig 4). Common type 1 fimbriae that mediate adhesion, including *fimA-I* and *ompA,* were present in *E. coli* isolates (S3 Fig). Other common virulence factors present in *E. coli* isolates include Yersiniabactin virulence factors *YbtA, YbtE, YbtP, YbtU, YbtX, YbtO,* enterotoxin senB, and aerobactin virulence. We identified many capsular polysaccharides in the *S. aureus* isolate, including (*cap8A-P*), iron-related proteins *isdA-G*, and enterotoxins *hlgA-C* and *hla/hly, hlb, hld*. Among the E. *faecalis* isolates, adhesins such as *efaA*, *ebpA-C*, and capsular polysaccharides *cpsB-I* were detected (S3 Fig).

## Discussion

In this study, we employed Nanopore Sequencing technology to comprehensively profile the antimicrobial resistance (AMR) mechanisms present in clinically resistant isolates. The application of Nanopore sequencing offers real-time, high-resolution genetic analysis that allowed us to uncover novel insights into the genetic basis of AMR in these isolates.

Based on the results, ST53 was the most common identified strain in *E. coli.* ST53 has been previously identified in *E. coli* isolates across different parts of the world that showed resistance to colistin [15, 16]. In *E. cloacae,* previously identified ST78, ST88 and ST98 were also observed in this study. ST78 is recognized as one of the global resistant clones of Extended-spectrum β-lactamase (ESBL) producing *Enterobacter cloacae* complex (ECC) and carbapenem-resistant *E. cloacae* complex (CREC). ST78 clones in the United States and Japan have been shown to harbor different carbapenemases genes highlighting their unique ability to acquire multidrug resistant plasmids [17, 18]. *S. haemolyticus* was assigned to ST30 and ST1 which have been detected in different clinical samples such as pus, blood and sputum [19]. ST30 found in this study has been previously implicated in vancomycin resistance [20]. Among the *A. baumannii* isolates, ST1 clone which previously has been predicted to show resistance against beta-lactams, fluroquinolones, aminoglycosides, sulfamethoxazole and carbapenem resistance was reported in the study [21]. Unfortunately, the sequence type of some isolates could not be assigned to any scheme due to the presence of single nucleotide substitutions and therefore the nearest ST were provided.

Our analysis revealed a diverse array of genetic elements associated with antimicrobial resistance. Notably, we identified a range of known resistance genes, including those encoding beta-lactamases, efflux pumps, and modifying enzymes. All *A. baumannii* isolates sequenced in this study harboured carbapenemase genes belonging to class D beta-lactamases. The carbapenem-hydrolyzing β-lactamases in *A. baumannii* are either metallo-β-lactamases (MBLs) or oxacillinases (carbapenem-hydrolyzing class D β-lactamases [CHDLs]). However, in this study, we didn’t detect any MBL’s in our study similar to another study in Morocco and Turkey(32-33). Intriguingly, we observed instances of co-occurring resistance mechanisms within individual isolates. Three out of five isolates of *A. baumannii* co-harboured carbapenemase and extended spectrum beta-lactamase genes. This occurrence of double or multiple carbapenemase genes in a single isolates is a finding demonstrated in other studies [22] as well as co-existence of carbapenemase and extended-Spectrum *β*-Lactamase genes in single isolate [24]. One gram-positive isolate harboured carbapenems hydrolyzing oxacillinase which has been described elsewhere [25]. Interestingly, all *E. coli* isolates (4/4) harbored more than three ESBL genes with one isolate demonstrating presence of 7 ESBL genes belonging to three classes of ESBLs (Class A, C, D). This phenomenon highlights the complexity of AMR and the potential for multiple mechanisms to synergistically contribute to resistance. Understanding these interactions is crucial for devising effective treatment strategies.

Besides beta-lactam resistance genes, we also found other genes which confer resistance to a wide variety of antibiotics including aminoglycosides, tetracycline, sulfonamides, chloramphenicol, macrolides, and fosfomycin. This points out to the existence of multidrug resistance isolates especially in the case of *A. baumannii*, *E. coli, E. cloacae* and *S. haemolyticus. A. baumannii* infections are commonly treated using different antibiotics including aminoglycosides, carbapenems, sulbactam, tigecycline, piperacillin/tazobactam, and polymyxins E and B. Already studies have shown increased resistance of *A. baumannii* to aminoglycosides and tetracyclines [26–28]. The *A. baumannii* isolates harbored aminoglycoside resistance genes (*aph(6)-Id* and *aph(3’’)-Ib* and tetracycline resistance genes (*tet(B)* and *tet(X),* a finding similar to previous studies [29–32]. In addition, an isolate contained the resistance gene *armA* which has previously been shown to show high level of resistance against several aminoglycosides including gemtamicin, tobramycin, and amikacin [33]. In addition, we identified macrolide resistance genes *mphE* and *msr(E)* which commonly associates with multidrug resistance in *A. baummanii* isolates [33]. Among the *E. coli*, aminoglycoside resistance genes identified included *aac(3)-IId, aac(3)-IIe*, *aac(6’)-Ib-D181Y, aac(6’)-Ib-cr5, aph(3’’)-Ib*, and *aph(6)-Id,* concordant with previous studies [26–28]. Resistance genes present in *E. coli* isolates against other commonly used inexpensive and readily available drugs such as quinolones, trimethoprim and sulfonamide and increase the risk of rendering these drugs ineffective in the long run. Furthermore, the emergence of methicillin resistant *S. aureus* and *S. haemolyticus* strains is particularly concerning and increases the risk of nosocomial and community acquired infections. The coexistence of multi-drug resistance underscores and compounds the challenge associated with treating infections caused by these pathogens.

Nanopore Sequencing enabled the detection of genetic elements such as mobile genetic elements (MGEs) and plasmids. In the present study a high proportion of clinical bacterial isolates from inpatients at Thika Level V hospital was found to carry plasmid replicons. The present findings are in concordance with previous studies elsewhere [34]. The observed high carriage of plasmid replicons by the analyzed isolates might plausibly be a reflection of resistance selection pressure due to high antibiotic exposure in hospital settings. Plasmids are infectious double stranded DNA molecules that are found within bacteria. Horizontal gene transfer promotes successful spread of different types of plasmids within or among bacteria species, making their detection an important task for guiding clinical treatment. Plasmid replicon type IncF predominantly identified in *E. coli* is frequently associated with genes encoding carbapenemases, aminoglycoside-modifying enzymes and plasmid-mediated quinolone resistance [35]. Various MDR plasmids predicted in this study associated with conjugative plasmid replicons IncFIB(K)_1_Kpn3, IncFIB(AP001918)_1 AP001918, IncI1_1_Alpha, and RepA_1_pKPC-CAV1321_CP011611. These plasmids are known to be associated with different several resistance profiles and carry both AMR and VF genes[36]. The largest number of plasmid types were found amongst *E. coli* strains. These strains acquire such as they are essential for pathogenicity[37]. Specific plasmids types such as ColRNAI_1 have been found in both *A. baumannii* and *E. coli* isolates and its implicated to carry the bla-CMY gene, which is an ampC type ESBL [38]. In terms of integrons, we found the presence of class 1 integrons in *E. cloacae, E. coli* and *A. baumannii* isolates. The AMR genes *blaOXA-10, QnrVC* and *arr* in *E. cloacae*, *blaOXA-1, catB, arr* and *qacE* in *E. coli* and *aar* in *A. baumannii* were encoded within an integron-integrase gene (Intl1). Class 1 integrons cassettes carry antibiotic resistance genes making them a significant player in the spread of AMR[39]. Our analysis uncovered the presence of diverse MGEs carrying resistance determinants, emphasizing their role in disseminating AMR genes within and between bacterial populations. These findings underscore the need for continued surveillance and control of MGE-mediated spread of resistance.

Apart from AMR genes, pathogenic bacteria have developed virulence factors against host defense mechanisms. In our study we identified and classified various different types of virulence factors (i.e immune evasion, adhesion, iron acquisition systems, surface coats, secretion systems enterotoxins). The presence of these virulence factors serves to increase the pathogenicity of different bacterial species and may also be spread to other bacterial organisms via horizontal gene transfer. Some key virulence genes identified in *E. coli* such as *fimA-I* and *ompA* which mediate adhesion. The expression of these surface adhesins increases the virulence of *E. coli* strains by ensuring close contact between the bacteria and host cell. In addition, Yersiniabactin virulence factors commonly associated with urinary tract infections were also identified in *E. coli* isolates[40]. In addition, the presence of senB enterotoxin implicated in the development of severe diarrhea among patient infected with enteroinvasive *E. coli* and *shigella* was also reported [41]. In the *S. aureus* isolate, capsular polysaccharide, *cap8A-P* identified. These capsular polysaccharides are covalently attached to peptidoglycan and play several roles including biofilm formation, colonization and evading phagocytes uptakes and protecting important bacterial cell wall constituents [42]. Other virulence factors such as enterotoxins *hlgA-C* is a cytolytic pore forming toxin that kills polymorphonuclear phagocytes, disrupt endothelial and epithelial barriers [41].

In conclusion, this study demonstrates the utility of Nanopore Sequencing in unraveling the intricate landscape of antimicrobial resistance in clinically resistant isolates. Our findings provide valuable genetic insights into the mechanisms underpinning resistance, including the detection of novel mutations and the role of MGEs. These results contribute to our broader understanding of AMR and have implications for guiding clinical management and public health efforts in combatting antimicrobial resistance. Future directions include expanding this approach to larger and more diverse sample sets to further explore the breadth of resistance mechanisms. Additionally, functional studies will be essential to validate the impact of newly identified mutations and better understand their clinical significance. As Nanopore Sequencing technology continues to evolve, it holds promise for continued advancements in our understanding of antimicrobial resistance and its implications for global health.

## Materials and Methods

### Ethics statement

This study was approved by the Institutional Scientific and Ethics Research Committee (ISERC) of Mount Kenya University (MKU/ERC/1687) and licensed by National Commission for Science, Technology, and Innovation (NACOSTI) (NACOSTI /P/21/7678). Before enrolment, informed consent was obtained from all participants or their legal guardian.

### Bacterial isolates and DNA extraction

Bacterial isolates (n=202) collected between March and November 2021 from patients at Thika Level V Hospital were retrieved from frozen growth in glycerol stocks from the MKU research laboratory. The bacterial isolates had been previously identified by Vitek Machine. Antimicrobial susceptibility testing was carried out using the Kirby-Bauer disk diffusion method. Isolates were tested against carbapenems (imipenem and meropenem), cephalosporins (cefuroxime, ceftriaxone, cefepime and ceftazidime, cefotaxime), cephamycins (cefoxitin), monoamides (aztreonam), quinolones (ciprofloxacin and levofloxacin), aminoglycosides (erythromycin, gentamicin and amikacin), beta-lactams (penicillin, cloxacillin, piperacillin, amoxicillin, and ampicillin), lincosamidess (clindamycin), glycopeptides (vancomycin) tetracyclines (Minocycline and tetracycline) and sulfonamides (sulfamethoxazole). Diffusion diameters were interpreted based on the Clinical Laboratory Standards and Institute (CLSI) guidelines as resistant (R), susceptible (S), or intermediate (I).

For genomic surveillance, 17 isolates which showed resistance against multiple antibiotics were selected for whole genomic sequencing (Supp A). The isolates were grown on nutrient agar for 18 hours at 37^0^C. After that, colonies were picked to create a culture suspension in 1X PBS, followed by genomic DNA extraction using Zymogen® bacterial and fungal DNA extraction kit. The quality and quantity of genomic DNA was then assessed using the NanoDrop and Qubit 3.0 fluorometer.

### ONT library preparation and sequencing

DNA library preparation was carried out using the SQK-LSK109 ligation sequencing kit. The fragmented DNA was first repaired using the NEBNext FFPE DNA Repair Mix and NEBNext Ultra II End Repair/dA-Tailing Module (New England BioLabs). Subsequently, individual barcodes were incorporated into the dA-tailed DNA using the EXP-NBD104 and EXP-NBD114 native barcoding expansion kit following the ONT protocol with NEB Blunt//TA Ligase Master Mix (New Englands Biolabs). Barcoded DNA samples were then equimolarly pooled, and adapters were attached using the Quick T4 DNA Ligase Quick Ligation Module (New Englands Biolabs). Before sequencing, the number of active pores on the flow cell R9.5(FLO-MIN106) was assessed, and equimolar pooling of samples was performed. Finally, sequencing was then carried out on the MinKNOW for 48 hours.

### Quality control

Basecalling of the raw Fast5 files produced after sequencing was performed using Guppy version 6.4.6 with the high accuracy mode option without quality filtering options. The multiple FASTQ files were merged into one and demultiplexed using Guppy barcoder [43]. FastQC was used to check the quality of the sequences. Afterward, Porechop (https://github.com/rrwick/Porechop) was used to trim adapters, while Filtlong (https://github.com/rrwick/Filtlong) filter reads with less than 500 base pairs. These steps contribute to the generation of high-quality sequences for downstream analysis.

### De novo assembly, polishing, and annotation

Flye. 2.9.1 (https://github.com/fenderglass/Flye) generated the draft assembly from the processed FASTQ reads [44]. The resulting draft assembly was then indexed and mapped against the individual reads using BWA-MEM (https://github.com/bwa-mem2/bwa-mem2) [45]. The generated SAM files were sorted, indexed, and converted to BAM format using Samtools (https://github.com/samtools/samtools). To improve the assembly quality, a three-step polishing approach was employed. Four rounds of Racon (https://github.com/isovic/racon) were first employed to polish the assembly using the mapped nanopore reads [46]. Subsequently, Medaka 1.7.2 (https://github.com/nanoporetech/medaka) and Homopolish were subsequently used to improve polih the resulting reads. The quality of the draft genome assembly was then assessed using Quast (https://github.com/ablab/quast), which provides comprehensive metrics including assembly accuracy and contig statistics [47]. Completeness of the draft assembly was evaluated using BUSCO (https://github.com/WenchaoLin/BUSCO-Mod), which checks the presence of evolutionarily conserved genes providing insights into genome completeness [48]. Genome completeness was expressed as BUSCO scores and included complete, fragmented, and missing BUSCOs, indicating /high-identity, partially present, and absent genes, respectively. Finally, the draft genome was annotated using Prokka (https://github.com/tseemann/prokka), which predicts coding sequences and other genomic features from bacterial genomes [49].

### Species identification

The multi-locus sequence typing (MLST) predicted bacterial species and sequence types in individual samples. Genomes were scanned against the traditional PubMLST typing schemes based on the existence of seven housekeeping genes. Default settings of the MLST scheme including 95% minimum identity of a complete allele, 10% as the minimum coverage of a partial allele, and 50 as the minimum score to match a scheme, were used.

### In-silico prediction of ARG, virulent genes, MGEs, plasmids, and integrons

Prediction of antimicrobial resistance genes was conducted using Abricate 1.0.1 (https://github.com/tseemann/abricate). The latest updated NCBI and VFDB databases were loaded in Abricate using the default setting and used to predict antimicrobial resistance and virulence genes[50]. Plasmids and integrons were predicted using Integron Finder and PlasmidFinder, respectively [51, 52]. AMR genes, virulence genes, and plasmids with >=90% sequence identity and >= 90% coverage and integrons with an e-value less than 0.0001 were analysed. The workflow from basecalling to identifying AMR genes is summarized below (S4 Fig).

## Acknowledgments

We appreciate the study volunteers and research teams from Thika Level V referral hospital, Kiambu County and the research team at Center for Malaria Elimination, Mount Kenya University for their technical assistance.

## Supporting information

S1 Fig. Genome completeness score using BUSCO before polishing.

S2 Fig. Genome completeness score using BUSCO after polishing.

Completeness score after four rounds of polishing using Racon followed by one round of Medaka and Homopolish.

S3 Fig. Virulence genes and classes present in *E. coli, E. faecalis,* and *S. aureus* isolates

S4 Fig. Flow diagram from reads pre-processing to the identification of AMR genes

S1 Table. Genome assembly summary statistics after multiple rounds of polishing.

S2 Table. MLST table revealing different organisms and sequence types (ST).

## Author Contributions

**Conceptualization:** Jesse Gitaka, Bernard N. Kanoi, Rachel Kimani, Sebastian Musundi

**Data Curation:** Sebastian Musundi, Rachel Kimani

**Formal Analysis:** Sebastian Musundi, Rachel Kimani, Patrick Wakaba

**Funding acquisition:** Jesse Gitaka, Bernard Kanoi

**Investigation**: Rachel Kimani, Sebastian Musundi, Patrick Wakaba, David Mbogo, Suliman Essuman, Bernard N. Kanoi, Jesse Gitaka

**Methodology:** Sebastian Musundi, Rachel Kimani, Patrick Wakaba, David Mbogo, Suliman Essuman, Bernard N. Kanoi, Jesse Gitaka

**Project Administration:** Jesse Gitaka

**Resources:** Jesse Gitaka, Bernard Kanoi

**Supervision:** Jesse Gitaka, Bernard N. Kanoi, David Mbogo, Suliman Essuman

**Validation:** Jesse Gitaka, Bernard N. Kanoi, Rachel Kimani, Sebastian Musundi

**Visualization:** Sebastian Musundi

**Writing - original draft:** Rachel Kimani, Sebastian Musundi, Patrick Wakaba

**Writing - review & editing:** Jesse Gitaka, Bernard N. Kanoi, David Mbogo, Suliman Essuman

## Notes

### Competing Interest Statement

The authors have declared no competing interest.

